# Non-invasive and painless mid-infrared modulation increases collagen in human and mouse skin

**DOI:** 10.1101/2024.03.28.587170

**Authors:** Zeyu Wang, Jiahui Zhu, Yuting Wang, Shuai Chen, Senlin Xu, Yaoying Li, Tianxing Hu, Yang Li, Xuanyue Wang, Renyue Ji, Sunny C. Li, Yan Yang, Hongbo Jia, Xiaowei Chen, Xing Fan, Lan Ge, Jianxiong Zhang

## Abstract

Stimulating collagen production in skin helps to enhance vitality while decelerating aging-associated processes in skin tissue. However, current approaches to enhancing collagen production are commonly limited by accompanying pain and trauma. Here, we report that mid-infrared modulation (MIMO) at an intensity of 70 mW/cm^2^ promotes collagen production in human or mouse skin in vivo without generating excessive heat. We found that protein levels of the collagen-degrading endopeptidase, MMP-1, were decreased in the skin of mice following MIMO treatment, whereas the collagen synthesis-related factors, TGF-β, HSP47, and HSP70, were all increased. In addition, MIMO stimulated collagen secretion in human dermal fibroblasts in vitro. This work demonstrates that MIMO is an effective, non-invasive, and painless intervention for in vivo enhancement of collagen production in the skin.

**One Sentence Summary:** Mid-infrared modulation promotes collagen production

## INTRODUCTION

As the largest organ (*1–4*), the skin performs a multitude of vital functions, such as safeguarding against physical trauma (*1, 2, 5*), regulating temperature and fluid balance (*6*), detecting painful or pleasurable environmental stimuli (*7*), and contributing to vitamin D synthesis (*8*). The dermis is a thick layer of fibrous and elastic tissue that confers flexibility and resilience to skin, while also housing nerve endings, sweat glands, sebaceous glands, hair follicles, and blood vessels (*9*). As the most abundant protein in the dermal extracellular matrix (ECM), collagen plays an essential role in cell adhesion and migration, as well as the regulation of cellular growth and metabolism (*2, 10–13*). Constituting approximately 75% of the dry weight of skin, collagen imparts both tensile strength and elasticity to this vital organ (*6, 13*).

However, various physiological and pathological factors can contribute to skin aging, resulting in the gradual emergence of a rough texture, wrinkles, spots, and atrophy, ultimately leading to the loss of skin tissue integrity (*1, 14–16*). In the process of natural skin aging, collagen fibers are destroyed and fragmented collagen fibers accumulate in the ECM (*17, 18*). The collagen may then undergo progressive cross-linking and calcification, decreasing skin elasticity (*19*). The ability of fibroblasts to synthesize collagen also gradually diminishes with age, leading to the deterioration and collapse of the ECM, and skin aging (*20, 21*). Skin aging is also accompanied by the activation of pro-inflammatory factors in skin tissues, and the resulting inflammatory response can further degrade collagen (*22, 23*). Additionally, cell senescence or apoptosis in skin tissue may directly disrupt the balance between tissue degradation and regeneration (*24, 25*). Photoaging, caused by long-term exposure to ultraviolet A (UVA) radiation from the sun, is the main cause of skin aging (*26*). UVA damages dermal fibroblasts by increasing cellular production of reactive oxygen species (ROS) (*27*), which affects ECM synthesis and remodeling in the dermis (*28*). When collagen, elastic fibers, and other dermal components decrease, the skin loses its elasticity, and wrinkles appear (*29*).

The irreversible senescence and the perpetual pursuit of a young and healthy appearance present a well-documented psychological paradox for humans (*30*). As a result, effectively repairing damaged skin and delaying aging are long-standing, primary objectives in the field of aesthetic medicine. To date, injection of exogenous collagen has been widely accepted as a non-surgical means for delaying skin aging. Recent studies have shown that exogenous collagen treatment can stimulate dermal fibroblasts to produce new collagen to repair skin tissues (*31, 32*). In addition, collagen is highly conserved in the evolution of different species, and thus exogenous collagen, such as that derived from bovine or porcine skin, is highly homologous to human collagen (*33, 34*), exhibiting low immunogenicity, high safety, and biocompatibility (*35*). In addition, some dermal fillers are also used to alleviate skin aging. Hyaluronic acid (HAc) filler has relatively high bioactivity and biocompatibility properties compared to other fillers(*36*). Crosslinked HAc filler stimulates fibroblast production of collagen fibers through the TGF-β/Smad pathway (*37–39*), although the precise pathway is still under debate. In recent years, oral collagen supplements have become popular as anti-aging products through increased marketing to consumers, in part because oral supplementation of collagen hydrolysates can reach deeper layers of the skin to rejuvenate skin physiology and appearance by enhancing hydration, elasticity, firmness, and reducing wrinkles (*40*).

Photoelectronic therapies are another widely accepted approach to alleviating skin aging that accounts for a substantial market share of medical aesthetics interventions for skin, such as radiofrequency (RF)(*41*). The mechanism of RF is through resistive heating within skin layers or subcutaneous tissues to transform RF into thermal energy (*42*). Molecular and histopathological studies have shown that RF results in the upregulation of cytokines and growth factors, remodeling, and reorienting collagen and elastin to increase the thickness of the papillary dermis (*43, 44*). These changes result in skin tightening and reducing scars and wrinkles (*44*). In addition to RF, near-infrared light devices have also been demonstrated to provide skin-tightening effects (*45*). Wavelengths in the infrared spectrum are absorbed by water in the skin, causing dermal heating (*46*). For example, a device irradiating at wavelengths in the 1100 to 1800 nm range was found to cause immediate skin tightening with effects lasting up to 3 months (*47*). However, the effects of mid-infrared light components of the infrared band, characterized by longer wavelengths and lower energy compared to the near-infrared band, remain inadequately investigated regarding their impact on skin. Here, in this study, we examined whether delivering mid-infrared modulation (MIMO) could significantly increase collagen thickness in the skin.

## RESULTS

### MIMO increases collagen in human skin

For mid-infrared light emission, we used a pulsable, thermal, near-blackbody infrared source to irradiate a 1 cm radius area of the forearm skin of 17 volunteers (9: 8, male: female). Subjects were exposed to MIMO for 10 min before undergoing ultrasonic measurement of collagen thickness (Fig. 1A). To ensure the safety of this test, subjects classified the heat-related effects of skin irradiation by MIMO on a scale of 0-4, with 4 representing maximum pain (Fig. 1B), and tests only proceeded if pain sensation of 2 or lower. Under illumination intensities ranging from 10 - 90 mW/cm^2^, all subjects experienced pain below 2 at 70 mW/cm^2^ (Fig. 1C), which led us to select this illumination level in subsequent tests. Preliminary skin collagen measurements showed that MIMO application to the forearm skin resulted in a significantly increased thickness of the collagen in all male and female subjects (Fig. 1D).

**Fig. 1.**
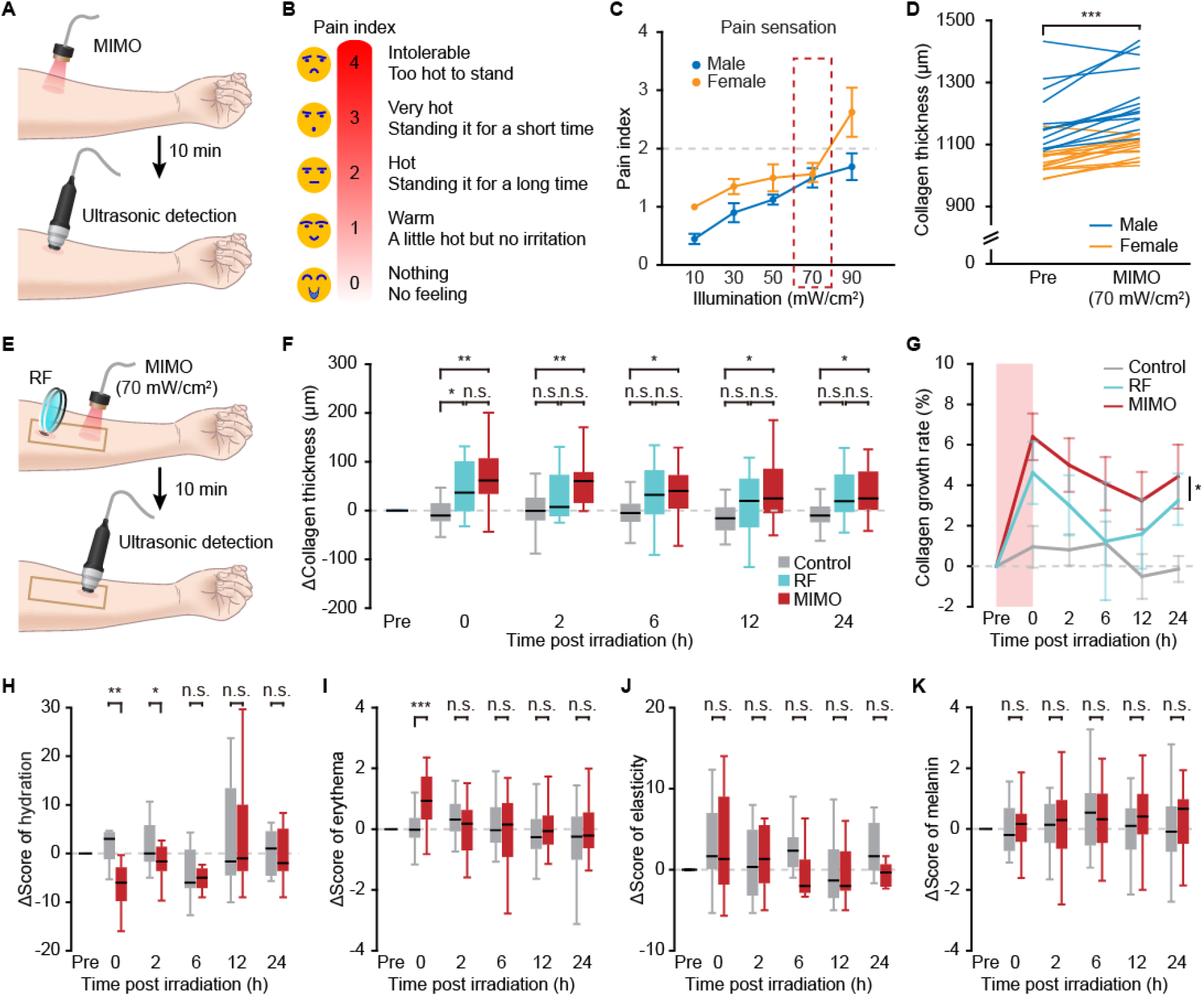
MIMO increases the thickness of collagen in human skin. (**A**) The experimental flow for MIMO irradiation on human skin. (**B**) Pain index, based on the pain sensation under MIMO application. (**C**) Pain sensation of male (n = 9) and female (n = 8) volunteers exposed to different MIMO illuminations (10 mW/cm², 30 mW/cm², 50 mW/cm², 70 mW/cm², 90 mW/cm²). Gray dashed line: pain index 2 (hot, a little hot but not irritation); Red dashed box: ideal MIMO illumination (70 mW/cm², selected for subsequent human experiments). (**D**) The changes in collagen thickness with 70 mW/cm²illumination in individual volunteers (27 pieces of skin from 17 volunteers (male 9, female 8)). *P* = 3.59e-05, Two-sided Wilcoxon sign-rank test. n.s. *p* > 0.05, \**p* < 0.05, \*\**p* < 0.01, \*\*\**p* < 0.001, applies for all figures. (**E**) Schematic for MIMO (70 mW/cm²) and RF experiments in human skin. (**F**) The changes in collagen thickness among MIMO (red) group, RF (blue) group, and control (gray) group within 24 h. *p*(0) _MIMO, Control_ = 0.0019, *p*(0) _RF, Control_ = 0.015, *p*(0) _MIMO, RF_ = 0.24, *p*(2) _MIMO, Control_ = 0.0038, *p*(2) _RF, Control_ = 0.15, *p*(2) _MIMO, RF_ = 0.26, *p*(6) _MIMO, Control_ = 0.048, *p*(6) _RF, Control_ = 0.083, *p*(6) _MIMO, RF_ = 0.71, *p*(12) _MIMO, Control_ = 0.039, *p*(12) _RF, Control_ = 0.14, *p*(12) _MIMO, RF_ = 0.60, *p*(24) _MIMO, Control_ = 0.021, *p*(24) _RF, Control_ = 0.051, *p*(24) _MIMO, RF_ = 0.74, Two-sided Wilcoxon sign-rank test. (**G**) The collagen growth rate for each time post irradiation in each group. Light red area: operation time, *p* (MIMO, RF) = 0.031, One-sided Wilcoxon sign-rank test. (**H**-**K**) The changes of other parameters of the skin, including hydration (**H**), *p*(0) = 0.0039, *p*(2) = 0.020, *p*(6) = 0.88, *p*(12) = 0.71, *p*(24) = 0.52; Erythema (**I**), *p*(0) = 5.76e -05, *p*(2) = 0.21, *p*(6) = 0.60, *p*(12) = 0.45, *p*(24) = 0.50; Elasticity (**J**), *p*(0) = 0.91, *p*(2) = 0.82, *p*(6) = 0.12, *p*(12) = 0.91, *p*(24) = 0.21; Melanin (**K**), *p*(0) = 0.21, *p*(2) = 0.54, *p*(6) = 0.72, *p*(12) = 0.11, *p*(24) = 0.22. Two-sided Wilcoxon rank-sum test.

To further assess collagen thickening in response to MIMO irradiation, we adopted a previously validated RF therapy as a positive control, while an untreated patch of forearm skin was measured as the negative control, with each subject receiving the treatment and both controls in spatially separated areas of forearm skin, and changes in collagen was monitored over the following 24 hours (Fig. 1E). Ultrasound measurements showed that the collagen thickened immediately following both RF and MIMO treatments, with the MIMO group showing stable, significantly thicker within 24 hours post irradiation (hpi) compared with the untreated control region. By contrast, the median collagen thickness of RF-treated skin trended higher than that of the untreated control, but did not reach statistical significance in the following detection time within 24 hpi (Fig. 1F). We also noted that although no obvious difference in collagen thickness was detected between the MIMO and RF groups during the observation period, the collagen growth rate (ΔThickness/ thickness untreated) was significantly higher and the thickening effect lasted longer in the MIMO group than the RF group (Fig. 1G). These findings suggest that MIMO produces more effective collagen thickening than RF.

In addition to collagen thickness, we also tested skin hydration, erythema, elasticity, and melanin contents before and after treatments. Among these parameters, skin hydration significantly decreased after MIMO irradiation, but returned to baseline at 6 hpi (Fig. 1H). In addition, erythema of the skin increased significantly after MIMO treatment, returning to baseline by 2 hpi (Fig. 1I). By contrast, neither skin elasticity nor melanin content exhibited any significant difference from controls within the 24h observation period (Fig. 1J and K).

### MIMO increases collagen in mouse skin

To further validate the effects of MIMO on enhancing collagen production in human skin, and to explore the most effective irradiation parameters, we adopted balb/c nude mice as an in vivo animal model due to their well-exposed skin. To assess pain sensation from different MIMO intensities, we evaluated pain responses (evasion or latency to paw withdrawal during treatment), body temperature, and damage to the skin following treatment applications (Fig. 2A). We applied MIMO illumination levels ranging from 10 - 130 mW/cm^2^ for 60 minutes to the flat surface of the mice back. If the mice displayed behaviors such as evading or retracting their paws during the experiment, we designated the duration of this time as pain response latency. The findings revealed that, if the illumination was more than 70 mW/cm^2^, the mice displayed obvious pain responses and had decreased latency to withdrawal (Fig. 2B). Skin temperature measurements showed a gradual increase along with irradiation intensity, stabilizing lower than 42 °C at 70 mW/cm^2^, which corresponded to a very mild level and did not exceed the threshold for heat-related pain channels(*48*) (Fig. 2C). Photographs captured during MIMO irradiation were used to assess changes or damage to skin (Fig. 2D). After irradiation (or upon escape of the mouse during irradiation, which was considered the end of the treatment), we observed that mice treated with MIMO intensity > 70 mW/cm^2^ displayed varying degrees of wounds to the skin surface, whereas no such damage was observed in mice treated with MIMO ≤ 70 mW/cm^2^, causing no damage to the skin surface compared to the control group (Fig. 2E).

**Fig. 2.**
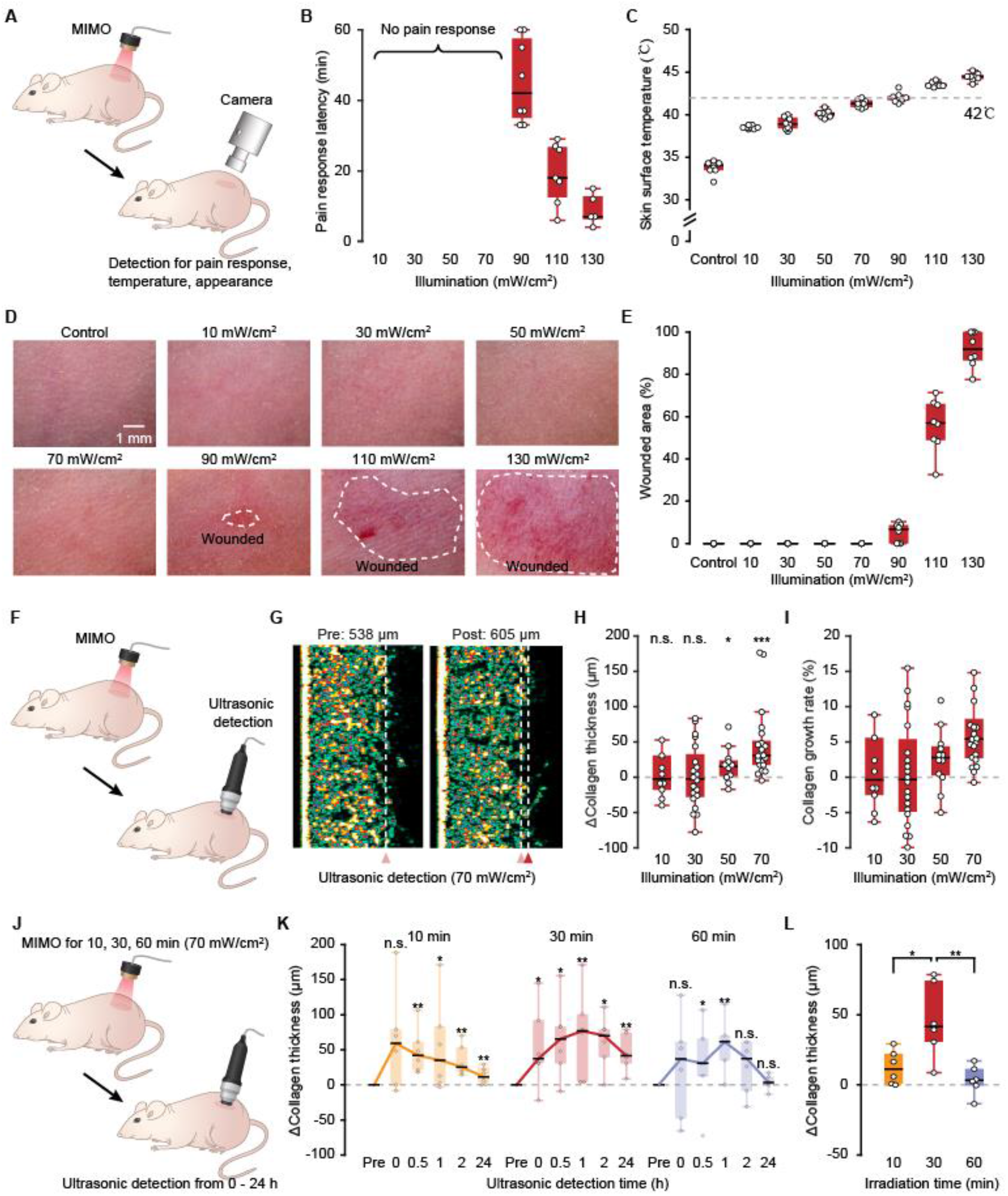
MIMO increases the thickness of collagen in mouse skin. (**A**) The experiment flow for safety evaluation of MIMO on mouse skin with gradient illumination, followed by detection for pain response, temperature, and appearance. (**B**) The pain response latency of mice within 60 min application of MIMO. (**C**) The temperature of skin surface within 60 min application of MIMO. (**D**) Appearances of mouse skin after MIMO application with different illuminations. The white dashed circle indicates the wounded area. (**E**) The wounded areas of mouse skin after MIMO application with different illuminations. (**F**) Schematic showing the general experimental flow. MIMO irradiates on the mouse skin for 10 minutes, followed by ultrasound detection of collagen thickness. (**G**) An example showing the comparison of collagen thickness in mouse skin before and after 70 mW/cm²MIMO application. (**H**) The changes in collagen thickness of mouse skin with different MIMO illuminations, *p*(10) = 0.86, n = 9; *p*(30) = 0.98, n = 20; *p*(50) = 0.019, n = 12; *p*(70) = 7.97e-5, n = 21, Two-sided Wilcoxon sign-rank test. (**I**) The collagen growth rate of mouse skin with different MIMO illuminations. (**J**) The experimental flow for MIMO effect with different irradiation times on mouse skin within 24 h. (**K**) The changes in collagen thickness with different irradiation times within 24 h (n = 6 mice), *p*(10 min)_0 h_ = 0.14, *p*(10 min)_0.5 h_ = 0.0022, *p*(10 min)_1 h_ = 0.048, *p*(10 min)_2 h_ = 0.0022, *p*(10 min)_24 h_ = 0.0022, *p*(30 min)_0 h_ = 0.015, *p*(30 min)_0.5 h_ = 0.048, *p*(30 min)_1 h_ = 0.0022, *p*(30 min)_2 h_ = 0.015, *p*(30 min)_24 h_ = 0.0022, *p*(60 min)_0 h_ = 0.36, *p*(60 min)_0.5 h_ = 0.048, *p*(60 min)_1 h_ = 0.0022, *p*(60 min)_2 h_ = 0.36, *p*(60 min)_24 h_ = 0.36, Two-sided Wilcoxon sign-rank test. (**L**) The increases in collagen thickness with different irradiation times at 24 h post irradiation, *p*(10 min vs 30 min) = 0.015, *p*(30 min vs 60 min) = 0.0087, *p*(10 min vs 60 min) = 0.31, Two-sided Wilcoxon rank-sum test.

Based on the above data, we next applied MIMO at 10, 30, 50, or 70 mW/cm^2^ for 10 min, and then detected the changes in the thickness of skin collagen by ultrasound (Fig. 2F). For example, when 70 mW/cm^2^ MIMO irradiated the skin of mice, the thickness of collagen increased from 538 μm to 605 μm (Fig. 2G). Although both 50 mW/cm^2^ and 70 mW/cm^2^ intensities resulted in significantly increased collagen thickening in skin (Fig. 2H), 70 mW/cm^2^ was associated with a higher growth rate of the collagen thickness compared to 50 mW/cm^2^ (Fig. 2I). As 70 mW/cm^2^ irradiation provided the greatest effects on collagen production without inducing pain or skin damage, we selected this treatment for further tests, which was consistent with the illuminance selection in human tests (Fig. 1C).

In subsequent tests with 24h observation periods, we assessed 70 mW/cm^2^ MIMO for 10 min, 30 min, and 60 min exposure times (Fig. 2J). Irradiation for 10 min, 30 min and 60 min caused varying degrees of collagen increase, but this effect was still maintained at 24 hpi only with 10 min and 30 min exposure (Fig. 2K), and the increased thickness of 30 min is higher than that of 10 min (Fig. 2L). Taken together, these results indicated that 70 mW/cm^2^ MIMO could increase collagen production in skin without apparent damage in mice and that exposure time could affect collagen thickness.

### MIMO does not upregulate pro-inflammatory factors and chemokine

To assess whether the effect of 70 mW/cm^2^ MIMO on collagen production was associated with pro-inflammatory response, we examined levels of inflammatory cytokines (i.e., TNF-α, IL2, IL6, IL12, and IL17) in the skin of treated and untreated mice at 24 hpi. ELISA-based detection indicated that no significant changes occurred in the production of any of these pro-inflammatory factors following MIMO exposure (Fig. 3A to F). We noted that levels of TNF-α, a hallmark of cancer (*49*), remained low following MIMO irradiation, suggesting that MIMO irradiation at appropriate levels does not induce skin damage. In addition, IL17 expression has been reported in senescent cells (*50*), and the stably low IL17 levels following MIMO irradiation suggest that MIMO does not contribute to skin aging. As the chemokine, CXCL1 is well established to be closely linked with the initiation and progression of skin disease (*51*). The expression of CXCL1 showed no significant difference before and after MIMO irradiation (Fig. 3G).

**Fig. 3.**
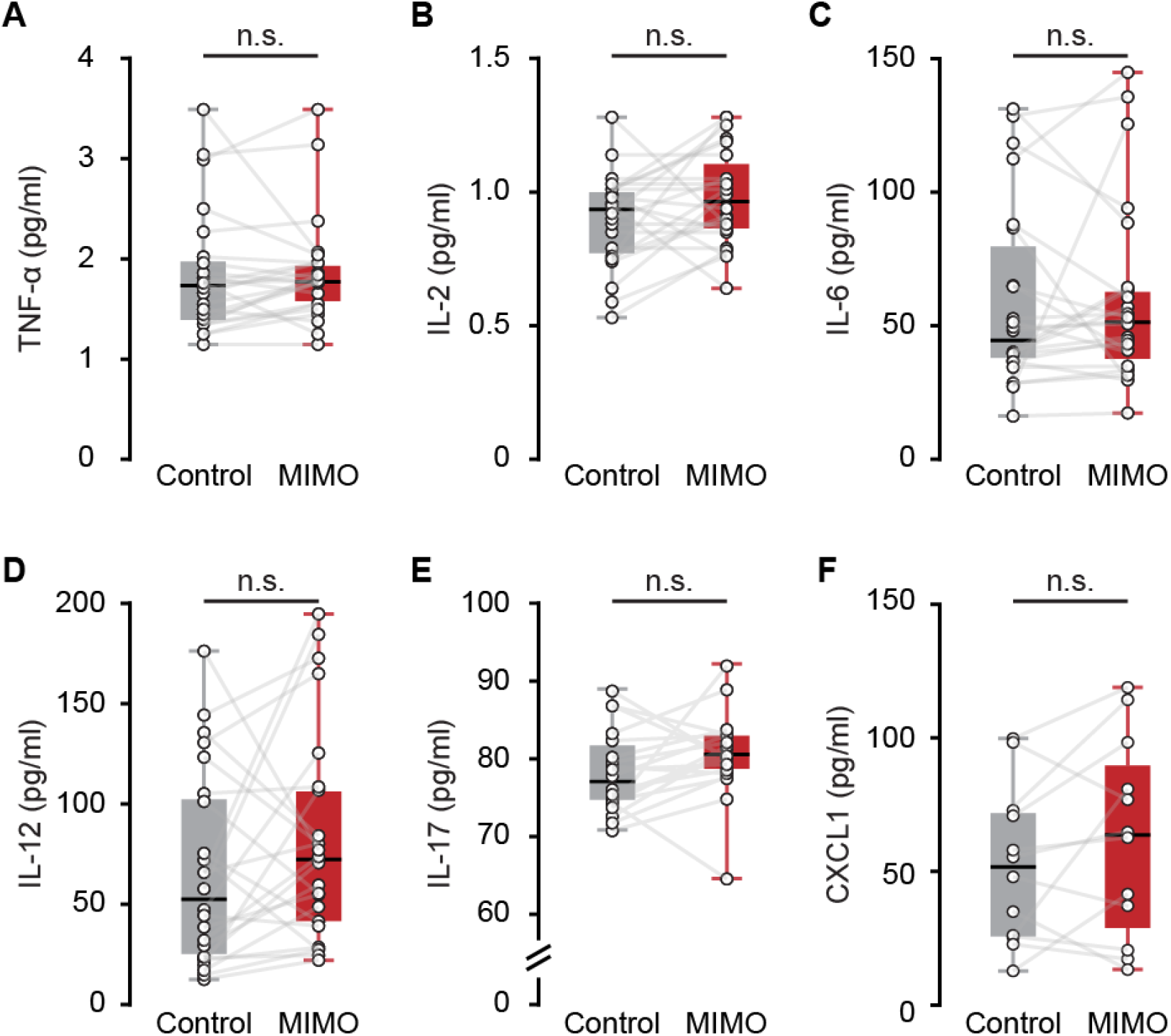
MIMO does not upregulate pro-inflammatory factors. (**A-E**) The changes of pro-inflammatory factors in mouse skin following 70 mW/cm²MIMO application, including TNF-α (**A**), n = 24, *p* = 0.17; IL-2 (**B**), n = 24, *p* = 0.09; IL-6 (**C**), n = 24, *p* = 0.75; IL-12 (**D**), n = 24, *p* = 0.19, and IL-17 (**E**), n = 20, *p* = 0.12, Two-sided Wilcoxon sign-rank test. (**F**) The changes of CXCL1 in mouse skin following 70 mW/cm²MIMO application, n = 12, *p* = 0.22, Two-sided Wilcoxon sign-rank test.

### Repeated MIMO irradiation prolongs the collagen thickening effect

To investigate the effects of repeated MIMO exposure on collagen production, we applied MIMO using the above parameters to the same area of skin in mice for 30 min per day for 7 consecutive days (Fig. 2L). Subsequently, we monitored changes in thickness of the skin collagen, using the contralateral skin of the same respective mice as the untreated controls (n = 13 mice; Fig. 4A). We observed that the collagen was significantly increased upon initial MIMO irradiation, and this effect was sustained for 7 consecutive days of irradiation. Continued monitoring post-treatment showed that the collagen thickening persisted for up to one month (Fig. 4B). After 7 consecutive days of irradiation, skin tissue samples were collected from the irradiated and control patches. Evaluation of collagen thickness by hematoxylin and eosin (H&E) staining (Fig. 4C) revealed that collagen had significantly increased in irradiated patches over in control samples (Fig. 4D), exhibiting a growth rate approaching 10% (Fig. 4E)

**Fig. 4.**
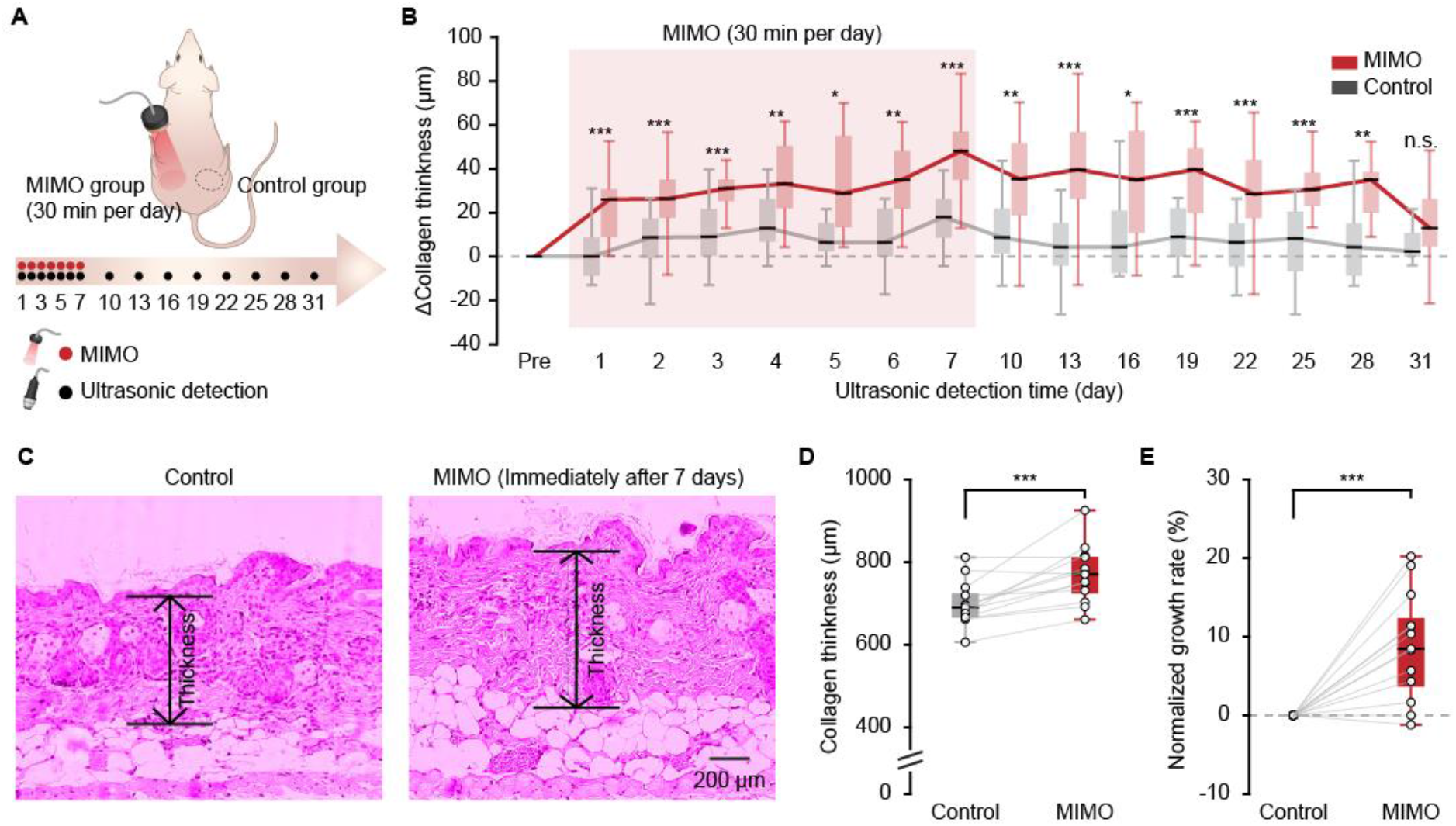
Repeated MIMO irradiation prolongs the collagen thickening effect. (**A**) The experimental flow for the MIMO effect over 31 days. Daily MIMO application (30 min per day) and daily ultrasound detection for the first 7 days, followed by ultrasound detection every 3 days for a total of up to 31 days. (**B**) The changes in collagen thickness between the MIMO (red) and control (gray) group over 31 days (21 mice). *p*(1) = 3.91e-04, *p*(2) = 2.13e-04, *p*(3) = 2.62e-04, *p*(4) = 0.0019, *p*(5) = 0.012, *p*(6) = 0.0034, *p*(7) = 1.32e-04, *p*(10) = 0.0057, *p*(13) = 6.81e-04, *p*(16) = 0.018, *p*(19) = 2.44e-04, *p*(22) = 2.13e-04, *p*(25) = 9.77e-04, *p*(28) = 0.002, *p*(31) = 0.065, Two-sided Wilcoxon sign-rank test. (**C**) Examples of H&E staining images of mouse skin (paraffin sections, sampled immediately after 7 days) in control and MIMO groups. Black arrows and lines indicate the thickness of dermal collagen. (**D**) The changes in measured collagen thickness of mouse skin sections (13 mice), *p* = 3.59e-05, Two-sided Wilcoxon sign-rank test. (**E**) The growth rate of collagen thickness of mouse skin slices (13 mice), *p* = 3.59e-05, Two-sided Wilcoxon sign-rank test.

### MIMO regulates collagen-related protein expression

To explore the underlying mechanism responsible for the observed, sustained increase in collagen following MIMO, we investigated changes in the expression proteins involved in collagen synthesis and secretion, including MMP-1, TGF-β, HSP47, and HSP70, at different time points during the experimental period based on protocols reported in above animal studies. For this analysis, the early stage was defined as the first 7 days following cessation of MIMO irradiation, while the late stage encompassed the subsequent period of up to one month. Skin samples were then collected from irradiated and control patches for ELISA-based quantification in both the early and late stages (Fig. 5A).

**Fig. 5.**
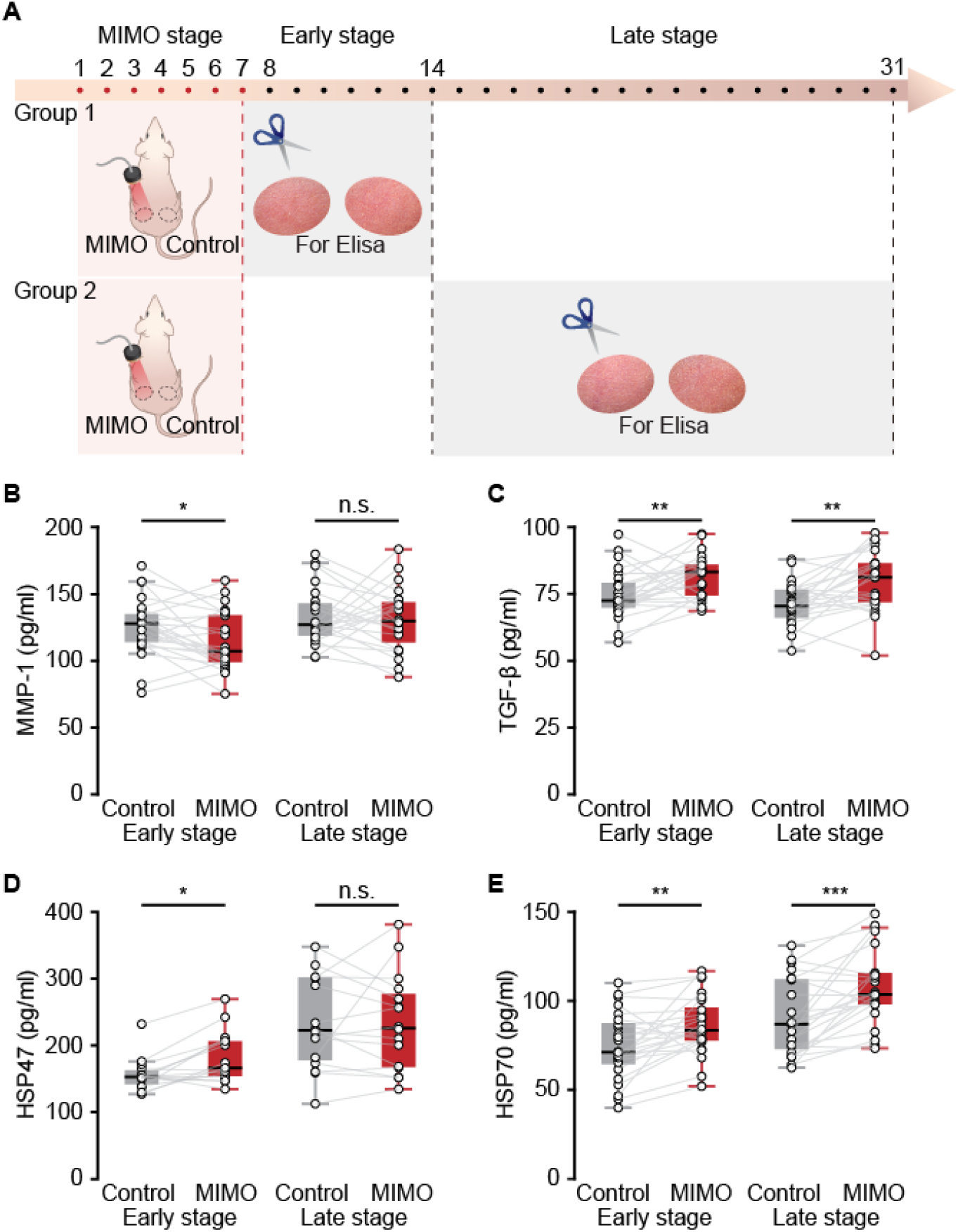
MIMO regulates the expression of collagen-related proteins. (**A**) Schematic showing the experimental flow: mouse skin was removed for Elisa in the two periods following the MIMO application. Arrow details three periods: the MIMO irradiation stage (1 - 7 days), the early detection stage (8 - 14 days), and the late detection stage (15 – 31 days). (**B-E**) The changes of collagen-related proteins in mouse skin of the MIMO and control groups in early and late detection stages, including MMP-1 (**B**), *p*(Early) = 0.020, n = 25; *p*(Late) = 0.41, n = 24; TGF-β (**C**), *p*(Early) = 0.008, n = 25; *p*(Late) = 0.0014, n = 24; HSP47 (**D**), *p*(Early) = 0.013, n = 15; *p*(Late) = 0.52, n = 24; HSP70 (**E**), *p*(Early) = 0.0012, n = 25; *p*(Late) = 6.07e-04, n = 24. Two-sided Wilcoxon sign-rank test.

Among these markers of collagen remodeling, we found that levels of the collagen-cleaving endopeptidase, MMP-1, were significantly decreased in MIMO-treated samples during the early stage (Fig. 5B). In contrast, TGF-β, which promotes extracellular matrix deposition and inhibits its degradation by MMPs (*52*), was significantly increased in both the early and late stages of the observation period in MIMO-treated skin (Fig. 5C). In addition, HSP47 was also upregulated in the early stage of MIMO treatment (Fig. 5D), while HSP70 was increased in the early and late stages (Fig. 5E), both of which proteins play a pivotal role in collagen synthesis and secretion (*53*).

These findings suggest that MIMO could potentially enhance collagen production while concurrently impeding collagen degradation.

### MIMO promotes collagen production in cultured human fibroblasts

Given the above evidence that MIMO may directly enhance collagen production, we next investigated whether and how this treatment affected human fibroblasts in vitro. To this end, we irradiated cultured human fibroblasts with 70 mW/cm^2^ MIMO for 10 min (Fig. 6A). Initial CCK8 cytotoxicity assays at 48 hours post MIMO treatment revealed that MIMO irradiation did not elicit any obvious decline in cell viability (Fig. 6B). Subsequent EdU assays examining the impact of MIMO on cellular proliferation showed that the number of cells expressing EdU significantly increased following MIMO irradiation (Fig. 6C and D). These findings suggest that MIMO irradiation could potentially enhance human fibroblast proliferation.

**Fig. 6.**
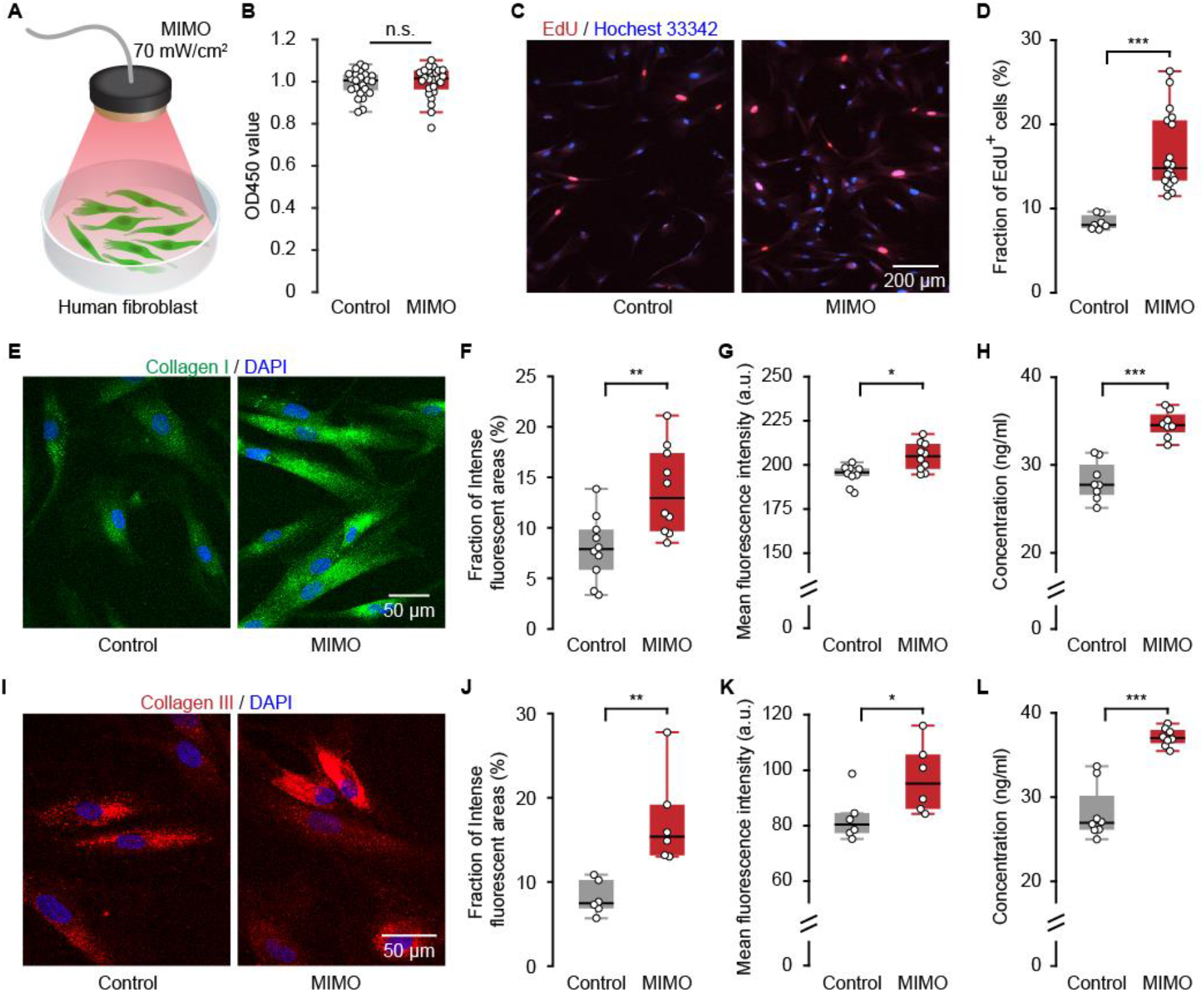
MIMO promotes collagen production in cultured human fibroblasts. (**A**) Cartoon showing the cultured HDFa with 70 mW/cm²MIMO application for 10 min. (**B**) Evaluation of radiation toxicity (indicated by the OD450 value) of MIMO. *p* = 0.43, n = 25. (**C**) Confocal images of stained HDFs in the control group and MIMO group. (**D**) The proportion of EdU+ cells in samples from control (n = 7) and MIMO group (n = 18). *p* = 1.6e-4. (**E**) Fluorescence images (collagen I stained green) of the control and MIMO group. (**F**) The proportion of intense fluorescent areas, *p* = 0.0058, n = 10. (**G**) The fluorescent intensity, *p* = 0.011, n = 10. (**H**) The concentration, *p* = 1.56e-4, n = 8. (**I**) Fluorescence images (collagen III stained red) of control and MIMO group. (**J**) The proportion of intense fluorescent areas, *p* = 0.0022, n = 6. (**K**) The fluorescent intensity, *p* = 0.026, n = 6. (**L**) The concentration, *p* = 1.56e-4, n = 8. Two-sided Wilcoxon rank-sum test.

As type I and III collagens formed at a higher proportion relative to other collagen types, and maintained in a fixed proportion in human skin (*54*), we directly assessed changes in these collagen types after MIMO irradiation. Immunofluorescent staining with antibodies targeting type I and type III collagen (Fig. 6E and I) showed that both the fraction of intense fluorescent areas defined as the top 20% of highest signal intensity regions, and the mean fluorescence intensity, significantly increased following MIMO irradiation compared with untreated controls (Fig. 6F and G for type I; Fig. 6J and K for type III). Subsequent quantification of collagen content by ELISA further indicated that both collagen types significantly increased in cells exposed to MIMO (Fig. 6H for type I, Fig. 6L for type III). These results suggest that MIMO irradiation directly promotes collagen production in human cells in vitro.

## DISCUSSION

In this study, we provide evidence that MIMO irradiation can be used as a painless and non-invasive treatment to increase collagen in human and mouse skin. Specifically, 70 mW/cm^2^ MIMO can increase the expression of proteins associated with collagen deposition, while inhibiting proteins associated with degradation, consequently stimulating a durable increase in skin collagen types I and III that can persist for one month after treatment for seven consecutive days, implying the deceleration of aging-associated collagen depletion. These findings suggest that 70 mW/cm^2^ MIMO holds the potential to enhance skin quality and provide innovative strategies for mitigating the aging process.

Non-surgical techniques for enhancing collagen to rejuvenate facial skin, such as exogenous dermal filling (*55*) and radiofrequency technology (*42*), are very popular due to their relative ease in achieving rapid improvements. Although considered safe, these procedures are also accompanied by some adverse reactions (*56*). As a component of normal skin, hyaluronic acid (*57*) injections generally do not cause allergic reactions, although isolated cases of granulomatous reactions are still reported (*58*). In addition, the use of RF treatments for skin rejuvenation is well-established and widely used due to their demonstrable efficacy in improving the skin (*59*). Common side effects primarily include erythema, swelling, and small crusts (*60*). However, a certain proportion of individuals may experience discomfort and pain during the procedure (*58*).

Compared to these methods, MIMO provides two main advantages as a novel technology for improving skin. First, this treatment is safe, painless and non-invasive, eliciting no nociceptive responses in either humans (Fig. 1C) or mice (Fig. 2B) at 70 mW/cm^2^ intensity, nor does it result in excessive elevation of skin surface temperature (Fig. 2C). It should be noted that ensuring safety relies on the careful selection of appropriate intensity levels, and MIMO can still potentially induce skin damage at levels above 70 mW/cm^2^ (Fig. 2D and E). Besides, the irradiation time is also a crucial parameter. Our findings demonstrate that MIMO irradiation for 60 minutes results in minimal collagen thickening at the 24-hour detection point, whereas the effect is less significant with 10 or 30 minutes of irradiation (Fig. 2K and L). This may be attributed to abnormal expression of elastin and fibrin in the skin due to prolonged heat exposure, which are subsequently degraded by various MMPs, such as MMP-12 (*61*). In addition to safety, MIMO also offers the advantage of high efficiency. Our results suggest that a single exposure to MIMO confers a significantly higher efficacy thickening effect on skin collagen than RF in the 24-hour period following treatment (Fig. 1G).

Numerous previous studies have explored the possible mechanisms leading to the biological effects of MIMO (*62*), which are likely relevant to its beneficial effects on the skin. Due to the differences of absorption spectra characteristics in different biological substances, tissue absorption of light irradiation in the mid-infrared band yields divers effects beyond a simple homogeneous heating. Biomarkers such as glucose, proteins, lipids, and urea exhibit strong but distinct absorption features in the mid-infrared spectrum (*63*). Therefore, we hypothesize that the observed increase in skin collagen induced by MIMO is the net effect influenced by multiple contributing factors. In addition, MIMO accelerates DNA unwinding (*64*), enhances the permeability of voltage-gated calcium channels (*65*) and potassium channels (*66*), increases neuronal excitability (*62*), modulate startle responses (*66*), and accelerate associative learning (*62*). Besides, MIMO has been shown to mitigate cognitive decline and modify gut microbiota in Alzheimer’s disease model mice (*67*). From molecules to cells, from animal behavior to degenerative diseases, MIMO has demonstrated stable regulatory effects, suggesting that MIMO may be a reliable, non-invasive approach to inducing potentially beneficial effects on humans.

Our study also has limitations, particularly the absence of long-term data from clinical trials. Nevertheless, we were able to confirm that MIMO has the potential to sustain collagen thickening for a duration of one month following a 7-day treatment in mice (Fig. 4B). The precise mechanisms underlying the increase in skin collagen due to MIMO need to be thoroughly investigated, as this is a crucial prerequisite for successfully translating the beneficial effects of MIMO from laboratory settings to clinical applications.

## MATERIALS AND METHODS

### Experimental Design

The objective of this study was to evaluate the collagen-enhancing effect of MIMO on fibroblasts in vitro, mouse skin in vivo, and human skin, while optimizing irradiation parameters and preliminarily investigating the mechanism underlying collagen increase by MIMO. Self-controlled designs were primarily employed for volunteers and experimental animals in the in vivo studies. Sample size determination was based on previous experience and statistical analysis, and no outliers were excluded from the study. For experiments involving statistical analysis, a minimum of six animals per group were used, with ’n’ representing the number of independent biological replicates indicated in the figure legend. To account for subtle differences between experimental and control groups, blind methods were maximally utilized during data collection and analysis. The sample size for each experiment reflects the number of independent biological replicates.

### Mid-infrared modulation (MIMO) and radiofrequency (RF) stimulation

MIMO was previously used to activated cortical neurons and enhance learning ability in mice (62). According the previous study, here, we used a high-power infrared transmitter as a mid-infrared light source. It was connected to a programmable DC Power Supply (RIGOL DP832) using insulated copper wire. For RF application, an RF device (frequency 50/60 Hz; output: 9 VDC, 2A; NEWA, P/N: ND-PRD00125, China) was employed as the control group in this study. The operation was strictly in accordance with the manufacturer’s directions.

### Subject

A total of 17 healthy subjects (9 males and 8 females) were recruited to undergo the treatment protocol. Inclusion criteria were: Age ranging from 18 to 40, healthy, with no previous history of skin diseases. Subjects who did not consent to the study or had any of the following criteria were excluded: female subjects of childbearing age who were pregnant, those who were lactating, or unwilling to take effective contraceptive measures; those with significant skin diseases in tested areas or with a previous history of severe skin diseases; those with a history of laser treatment, chemical exfoliation, soft tissue packing, injection of botulinum toxin type A, or other unknown substances in the previous six months; those patients with skin rejuvenation surgery history; those having undergone radiotherapy or other treatments interfering with healing; a history of skin allergy; or any medical condition the investigators deemed inappropriate for clinical trial participants. The Ethical Review Board of the First Affiliated Hospital of Third Military Medical University approved the study (Approval (A) KY2024010).

### Safety evaluation of human skin

We established the illumination gradient of the mid-infrared emitter (10, 30, 50, 70, and 90 mW/cm^2^) and conducted safety evaluations (characterized by pain index) in response to different irradiation parameters in human subjects. A pain quantification table (pain index 0 to 4, based on pain perception under MIMO application) was intended to quantify the pain perception of subjects under varied illumination. A pain scale of 0-4 corresponds to Nothing (no feeling), Warm (a little hot but without any irradiation), Hot (standing for a long time), Very hot (standing for a short time), and Intolerable (too hot to stand). To eliminate accidents, a pain index below 2 was defined as the critical index for accident safety.

### Collagen detection

Collagen thickness and ultrasound images were acquired using an ultrasound detector (Serial No.CO8440.05-248, DermaLab Combo, Cortex Technology ApS). The skin tested in this study was healthy, smooth, relatively flat, hairless, and free of wounds or scars. Therefore, we chose the inner skin of the human forearm for human experiments and flat back skin in mouse experiments. Additionally, the above experiments were performed in a non-closed normal environment (an operating room with a temperature of 20 ℃ and relative humidity of 50%-60%).

### Hydration, erythema, melanin, and elasticity detection

Hydration, Erythema, and melanin were measured by detectors (HYDRATION PIN: SERIAL NO.CO6440.01-165, SKIN COLOR: SERIAL NO.CO9440.01-165, Dermalab Combo, CORTEX TECHNOLOGY ApS), elasticity was detected by Skin Analysis Machine (CBS-802, DIGITAL TEST) in this study.

### Mouse

BALB/c-nu SPF-grade male mice (6 to 8 weeks old) were obtained from Sbev (Beijing) Biotechnology Co., LTD. The mice were kept in conditions corresponding to a cycle of 12 hours of light/12 hours of dark (darkness beginning at 19:00), with pathogen-free circumstances. The animal facility was maintained at temperatures from 20 to 24 °C while the humidity was maintained between 45 % and 65 %. The animals were maintained in a consistent environment for five days. All experimental practices were aligned with the institutional animal welfare guidelines and approval from the Third Military Medical University Animal Care and Use Committee. Mice possessing marks of skin damage or inflammation (dermatitis, wounds, or redness) were removed from studies to limit interfering with results.

### Safety evaluation of mouse skin

Animal skin safety evaluation used an illumination gradient of MIMO (10, 30, 50, 70, 90, 110, and 130 mW/cm^2^) and assessed the safety on animal skin (characterized by pain response, temperature, and appearance) based on various irradiation parameters. Pain response latency was utilized to assess the thermal cumulative pain response of animals to MIMO stimulation in our study. This response was defined as the time necessary to observe paw retraction, body writhing, and active escape during a specific irradiation period (60 minutes) under slight anesthesia (general 0.8%; leg flexion and paw retraction occurred when the toes of mice were pinched using tweezers). When the mice exhibited the above reactions, the experiment was immediately ceased (cutoff time 60 minutes). Skin surface temperature was identified using an infrared temperature detector (325 Pro, Thermal Intelligence, FOTRIC, Dongguan Anyfine Electronic Technology Co., Ltd). A camera characterized images of the appearance of irradiated mouse skin.

### Histology

Samples were fixed using neutral buffered formalin (10%) at room temperature, dehydrated, and embedded in paraffin blocks for histological assessment. Formalin-fixed, paraffin-embedded blocks were sectioned into 2 to 4 μm sections. H&E staining was employed to examine alterations in skin histological structure and dermal thickness. Images (20×) of three representative sites in each tissue section were obtained using a microscope (Leica, SP5, Germany), and three representative measurements per image were acquired. The measurement sites were chosen to be an appropriate width representing the overall thickness of the stained dermal collagen within the image, and the widest or narrowest parts were excluded. The mean value of a total of nine measurements was calculated and employed as the final dermal thickness.

### ELISA

The levels of MMP-1, TGF-β, IL-2, IL-6, IL-12, TNF-α, HSP47, HSP70, CXCL1, collagen I, and collagen III, were examined using ELISA kits purchased from BioChannel Biological Technology Co., Ltd., according to the manufacturer’s directions.

### Cell culture

The human fibroblast cell line, HDFa, was obtained from FuHeng Cell Center (Shanghai, China). The cells were grown using 1640 culture medium (Gibco, Carlsbad, CA, USA), including 10% fetal bovine serum (Gibco, Carlsbad, CA, USA) as well as 1x Penicillin-Streptomycin Solution within a humidified incubator at 37 ℃ and 5% CO_2_.

### Cell viability analysis

The HDFa cells were grown with 2x10^4^ cells per well within 96-well plates and cultured using a complete medium for 48 hours. MIMO (70 mW/cm^2^) irradiation of the cells was performed for 10 minutes, while unirradiated cells were employed as a control group. Following irradiation for 48 hours, 10 μL of Cell Counting Kit-8 reagent was placed into all wells and maintained at 37 ℃ for 4 hours. The absorption level at 450 nm was identified using a microplate reader.

### 5-ethynyl-2’-deoxyuridine (EdU) cell proliferation measurement

The propagation of HDFa cells was characterized using tagging of freshly produced DNA according to the integration of the thymidine analog, EdU, during DNA production. HDFa cells were grown in 24-well plates with cell crawls, and cells were stimulated through the inclusion of appropriate drugs to identify EdU-stained cell crawls based on the manufacturer’s directions for the BeyoClick™ EdU Cell Proliferation Kit using Alexa Fluor 594 kit (C0078S, Beyotime, China). A fluorescence microscope (Leica, SP5, Germany) was utilized to characterize the fluorescence images of EdU-stained cells as well as Hoechst 33342-stained nuclei.

### Immunofluorescence staining

HDFa cells were incubated with 2x10^4^ cells per well in 24-well plates and cultured using a complete medium for 48 hours. The cells were fixed using 4% paraformaldehyde for 10 minutes and permeabilized with 0.5% Triton X-100 for 5 minutes. Following rinses with TBST, cells were blocked using blocking buffer (5% goat serum in 1X PBS) for 1 hour, maintained with the denoted antibodies at 37 °C for 2 hours, and stained using DAPI (Sigma-Aldrich, St. Louis, MO, USA). Images were obtained using a fluorescence microscope (Leica, SP5, Germany).

### Statistics

To compare data between groups, we used the nonparametric Wilcoxon rank sum test (unpaired), and Wilcoxon signed-rank test (paired) to determine statistical significance (P < 0.05) between them. In the figures, the data presented in the box-and-whisker plot indicate the median (center line), 25th and 75th percentiles (Q1 and Q3), i.e., IQR (box), Q1-1.5 ×IQR and Q3 + 1.5 ×IQR (whiskers).

## Acknowledgments

The authors are grateful to Ms. Jia Lou for her help in composing and layout editing the figures.

## Author contributions

JX.Z., X.C. and L.G. conceived the project. JX.Z. and X.C. designed the experiments; Z.W., JH.Z, Y.W., S.X., S.C., Y. L., T.H., Y.L., and R.J. performed the experiments; Z.W., JH.Z, Y.W. and JX.Z. performed the data analysis; L.G., X.F., Y.Y., S.L., and X.C. inspected the data and evaluated the findings; L.G., X.F., X.C., and JX.Z. wrote the manuscript with the help of all authors. All authors read and commented on the manuscript.

## Data and materials availability

All data are available in the main text.

## References

1. C. W. Cao, Z. C. Xiao, H. Q. Tong, Y. T. Liu, Y. L. Wu, C. R. Ge, Oral Intake of Chicken Bone Collagen Peptides Anti-Skin Aging in Mice by Regulating Collagen Degradation and Synthesis, Inhibiting Inflammation and Activating Lysosomes. Nutrients 14, (2022).

2. E. S. Chambers, M. Vukmanovic-Stejic, Skin barrier immunity and ageing. Immunology 160, 116–125 (2020).

3. F. O. Nestle, P. Di Meglio, J. Z. Qin, B. J. Nickoloff, Skin immune sentinels in health and disease. Nat Rev Immunol 9, 679–691 (2009).

4. C. Blanpain, E. Fuchs, Epidermal stem cells of the skin. Annu Rev Cell Dev Bi 22, 339–373 (2006).

5. P. Pittayapruek, J. Meephansan, O. Prapapan, M. Komine, M. Ohtsuki, Role of Matrix Metalloproteinases in Photoaging and Photocarcinogenesis. Int J Mol Sci 17, (2016).

6. J. W. Shin, S. H. Kwon, J. Y. Choi, J. I. Na, C. H. Huh, H. R. Choi, K. C. Park, Molecular Mechanisms of Dermal Aging and Antiaging Approaches. Int J Mol Sci 20, (2019).

7. M. Schmelz, Neuronal sensitivity of the skin. Eur J Dermatol 21, 43–47 (2011).

8. E. Kechichian, K. Ezzedine, Vitamin D and the Skin: An Update for Dermatologists. Am J Clin Dermatol 19, 223–235 (2018).

9. M. J. Mienaltowski, N. L. Gonzales, J. M. Beall, M. Y. Pechanec, Basic Structure, Physiology, and Biochemistry of Connective Tissues and Extracellular Matrix Collagens. Adv Exp Med Biol 1348, 5–43 (2021).

10. K. E. Kadler, C. Baldock, J. Bella, R. P. Boot-Handford, Collagens at a glance. J Cell Sci 120, 1955–1958 (2007).

11. S. Chattopadhyay, R. T. Raines, Collagen-Based Biomaterials for Wound Healing. Biopolymers 101, 821–833 (2014).

12. M. J. Mienaltowski, D. E. Birk, Structure, Physiology, and Biochemistry of Collagens. *Progress in Heritable Soft Connective Tissue Diseases* **802**, 5–29 (2014).

13. M. D. Shoulders, R. T. Raines, Collagen Structure and Stability. *Annu Rev Biochem* **78**, 929–958 (2009).

14. M. Majidian, H. Kolli, R. L. Moy, Management of skin thinning and aging: review of therapies for neocollagenesis; hormones and energy devices. Int J Dermatol 60, 1481–1487 (2021).

15. S. Natari, K. E. Kim, S. I. Ryu, J. H. Park, I. H. Kim, Device-Induced Neocollagenesis: Profibrotic Response or True Neocollagenesis? Laser Surg Med 52, 1010–1019 (2020).

16. D. Orioli, E. Dellambra, Epigenetic Regulation of Skin Cells in Natural Aging and Premature Aging Diseases. Cells-Basel 7, (2018).

17. J. Varani, L. Schuger, M. K. Dame, C. Leonard, S. E. G. Fligiel, S. Kang, G. J. Fisher, J. J. Voorhees, Reduced fibroblast interaction with intact collagen as a mechanism for depressed collagen synthesis in photodamaged skin. J Invest Dermatol 122, 1471–1479 (2004).

18. C. C. Zouboulis, E. Makrantonaki, G. Nikotakis, When the skin is in the center of interest: An aging issue. Clin Dermatol 37, 296–305 (2019).

19. M. A. Cole, T. H. Quan, J. J. Voorhees, G. J. Fisher, Extracellular matrix regulation of fibroblast function: redefining our perspective on skin aging. J Cell Commun Signal 12, 35–43 (2018).

20. G. J. Fisher, J. Varani, J. J. Voorhees, Looking older - Fibroblast collapse and therapeutic implications. Arch Dermatol 144, 666–672 (2008).

21. M. Q. Man, S. J. Xin, S. P. Song, S. Y. Cho, X. J. Zhang, C. X. Tu, K. R. Feingold, P. M. Elias, Variation of Skin Surface pH, Sebum Content and Stratum Corneum Hydration with Age and Gender in a Large Chinese Population. Skin Pharmacol Phys 22, 190–199 (2009).

22. P. Meyer, P. Maity, A. Burkovski, J. Schwab, C. Müssel, K. Singh, F. F. Ferreira, L. Krug, H. J. Maier, M. Wlaschek, T. Wirth, H. A. Kestler, K. Scharffetter-Kochanek, A model of the onset of the senescence associated secretory phenotype after DNA damage induced senescence. Plos Comput Biol 13, (2017).

23. F. P. Beserra, L. F. S. Gushiken, A. J. Vieira, D. A. Bérgamo, P. L. Bérgamo, M. O. de Souza, C. A. Hussni, R. K. Takahira, R. H. Nóbrega, E. R. M. Martinez, C. J. Jackson, G. L. D. Maia, A. L. Rozza, C. H. Pellizzon, From Inflammation to Cutaneous Repair: Topical Application of Lupeol Improves Skin Wound Healing in Rats by Modulating the Cytokine Levels, NF-κB, Ki-67, Growth Factor Expression, and Distribution of Collagen Fibers. Int J Mol Sci 21, (2020).

24. J. Campisi, P. Kapahi, G. J. Lithgow, S. Melov, J. C. Newman, E. Verdin, From discoveries in ageing research to therapeutics for healthy ageing. Nature 571, 183–192 (2019).

25. A. S. Wang, O. Dreesen, Biomarkers of Cellular Senescence and Skin Aging. Front Genet 9, (2018).

26. J. Liu, W. Zhang, The Influence of the Environment and Clothing on Human Exposure to Ultraviolet Light. Plos One 10, (2015).

27. S. Kowalski, J. Karska, M. Tota, K. Skinderowicz, J. Kulbacka, M. Drag-Zalesinska, Natural Compounds in Non-Melanoma Skin Cancer: Prevention and Treatment. Molecules 29, (2024).

28. W. Xia, T. H. Quan, C. Hammerberg, J. J. Voorhees, G. J. Fisher, A mouse model of skin aging: Fragmentation of dermal collagen fibrils and reduced fibroblast spreading due to expression of human matrix metalloproteinase-1. J Dermatol Sci 78, 79–82 (2015).

29. Y. P. Gu, J. X. Han, C. P. Jiang, Y. Zhang, Biomarkers, oxidative stress and autophagy in skin aging. Ageing Res Rev 59, (2020).

30. S. Q. Hu, Z. H. Li, J. Cores, K. Huang, T. Su, P. U. Dinh, K. Cheng, Needle-Free Injection of Exosomes Derived from Human Dermal Fibroblast Spheroids Ameliorates Skin Photoaging. Acs Nano 13, 11273–11282 (2019).

31. J. C. Yu, J. Q. Wang, Y. Q. Zhang, G. J. Chen, W. W. Mao, Y. Q. Ye, A. R. Kahkoska, J. B. Buse, R. Langer, Z. Gu, Glucose-responsive insulin patch for the regulation of blood glucose in mice and minipigs. Nat Biomed Eng 4, 499–506 (2020).

32. K. Matsuo, H. Okamoto, Y. Kawai, Y. S. Quan, F. Kamiyama, S. Hirobe, N. Okada, S. Nakagawa, Vaccine efficacy of transcutaneous immunization with amyloid β using a dissolving microneedle array in a mouse model of Alzheimer’s disease. J Neuroimmunol 266, 1–11 (2014).

33. N. Modepalli, H. N. Shivakumar, M. T. C. McCrudden, R. F. Donnelly, A. Banga, S. N. Murthy, Transdermal Delivery of Iron Using Soluble Microneedles: Dermal Kinetics and Safety. J Pharm Sci-Us 105, 1196–1200 (2016).

34. Y. C. Kim, J. H. Park, M. R. Prausnitz, Microneedles for drug and vaccine delivery. Adv Drug Deliver Rev 64, 1547–1568 (2012).

35. M. Kim, H. Yang, H. Kim, H. Jung, H. Jung, Novel cosmetic patches for wrinkle improvement: retinyl retinoate- and ascorbic acid-loaded dissolving microneedles. Int J Cosmetic Sci 36, 207–212 (2014).

36. K. L. Beasley, M. A. Weiss, R. A. Weiss, Hyaluronic Acid Fillers: A Comprehensive Review. Facial Plast Surg 25, 86–94 (2009).

37. T. Iannitti, J. C. Morales-Medina, A. Coacci, B. Palmieri, Experimental and Clinical Efficacy of Two Hyaluronic Acid-based Compounds of Different Cross-Linkage and Composition in the Rejuvenation of the Skin. Pharm Res-Dordr 33, 2879–2890 (2016).

38. S. Paliwal, S. Fagien, X. J. Sun, T. Holt, T. Kim, C. K. Hee, D. Van Epps, D. J. Messina, Skin Extracellular Matrix Stimulation following Injection of a Hyaluronic Acid-Based Dermal Filler in a Rat Model. Plast Reconstr Surg 134, 1224–1233 (2014).

39. T. H. Quan, F. Wang, Y. Shao, L. Rittié, W. Xia, J. S. Orringer, J. J. Voorhees, G. J. Fisher, Enhancing Structural Support of the Dermal Microenvironment Activates Fibroblasts, Endothelial Cells, and Keratinocytes in Aged Human Skin. J Invest Dermatol 133, 658–667 (2013).

40. L. Bolke, G. Schlippe, J. Gerss, W. Voss, A Collagen Supplement Improves Skin Hydration, Elasticity, Roughness, and Density: Results of a Randomized, Placebo-Controlled, Blind Study. Nutrients 11, (2019).

41. D. McDaniel, R. Weiss, M. Weiss, C. Mazur, C. Griffin, Two-Treatment Protocol for Skin Laxity Using 90-Watt Dynamic Monopolar Radiofrequency Device With Real-Time Impedance Intelligence Monitoring. J Drugs Dermatol 13, 1112–1117 (2014).

42. M. El-Domyati, T. S. El-Ammawi, W. Medhat, O. Moawad, D. Brennan, M. G. Mahoney, J. Uitto, Radiofrequency facial rejuvenation: Evidence-based effect. J Am Acad Dermatol 64, 524–535 (2011).

43. M. El-Domyati, T. S. El-Ammawi, W. Medhat, O. Moawad, M. G. Mahoney, J. Uitto, Expression of transforming growth factor-β after different non-invasive facial rejuvenation modalities. Int J Dermatol 54, 396–404 (2015).

44. D. H. Suh, J. H. Choi, S. J. Lee, K. H. Jeong, K. Y. Song, M. K. Shin, Comparative histometric analysis of the effects of high-intensity focused ultrasound and radiofrequency on skin. J Cosmet Laser Ther 17, 230–236 (2015).

45. C. C. S. Martignago, C. R. Tim, L. Assis, V. R. Da Silva, E. C. B. Dos Santos, F. N. Vieira, N. A. Parizotto, R. E. Liebano, Effects of red and near-infrared LED light therapy on full-thickness skin graft in rats. Laser Med Sci 35, 157–164 (2020).

46. A. Gupta, T. H. Dai, M. R. Hamblin, Effect of red and near-infrared wavelengths on low-level laser (light) therapy-induced healing of partial-thickness dermal abrasion in mice. Laser Med Sci 29, 257–265 (2014).

47. Y. Tanaka, K. Matsuo, S. Yuzuriha, Long-Term Evaluation of Collagen and Elastin Following Infrared (1100 to 1800 nm) Irradiation. J Drugs Dermatol 8, 708–712 (2009).

48. D. R. Loyd, P. B. Chen, K. M. Hargreaves, Anti-Hyperalgesic Effects of Anti-Serotonergic Compounds on Serotonin- and Capsaicin-Evoked Thermal Hyperalgesia in the Rat. Neuroscience 203, 207–215 (2012).

49. F. Balkwill, TNF-α in promotion and progression of cancer. Cancer Metast Rev 25, 409–416 (2006).

50. P. Solá, E. Mereu, J. Bonjoch, M. Casado-Peláez, N. Prats, M. Aguilera, O. Reina, E. Blanco, M. Esteller, L. Di Croce, H. Heyn, G. Solanas, S. A. Benitah, Targeting lymphoid-derived IL-17 signaling to delay skin aging. Nature Aging 3, 688-+ (2023).

51. N. Wang, W. P. Liu, Y. F. Zheng, S. Q. Wang, B. W. Yang, M. Li, J. X. Song, F. X. Zhang, X. T. Zhang, Q. Wang, Z. Y. Wang, CXCL1 derived from tumor-associated macrophages promotes breast cancer metastasis via activating NF-κB/SOX4 signaling. Cell Death Dis 9, (2018).

52. R. Wisdom, L. Huynh, D. Hsia, S. Kim, RAS and TGF-β exert antagonistic effects on extracellular matrix gene expression and fibroblast transformation. Oncogene 24, 7043–7054 (2005).

53. E. E. Abd El-Fattah, A. Y. Zakaria, Targeting HSP47 and HSP70: promising therapeutic approaches in liver fibrosis management. J Transl Med 20, (2022).

54. R. Gref, C. Deloménie, A. Maksimenko, E. Gouadon, G. Percoco, E. Lati, D. Desmaële, F. Zouhiri, P. Couvreur, Vitamin C-squalene bioconjugate promotes epidermal thickening and collagen production in human skin. Sci Rep-Uk 10, (2020).

55. H. Q. Lv, N. Gao, Q. X. Zhou, Y. Wang, G. X. Ling, P. Zhang, Collagen-Based Dissolving Microneedles with Flexible Pedestals: A Transdermal Delivery System for Both Anti-Aging and Skin Diseases. Adv Healthc Mater 12, (2023).

56. L. Requena, C. Requena, L. Christensen, U. S. Zimmermann, H. Kutzner, L. Cerroni, Adverse reactions to injectable soft tissue fillers. J Am Acad Dermatol 64, 1–34 (2011).

57. S. N. A. Bukhari, N. L. Roswandi, M. Waqas, H. Habib, F. Hussain, S. Khan, M. Sohail, N. A. Ramli, H. E. Thu, Z. Hussain, Hyaluronic acid, a promising skin rejuvenating biomedicine: A review of recent updates and pre-clinical and clinical investigations on cosmetic and nutricosmetic effects. Int J Biol Macromol 120, 1682–1695 (2018).

58. S. Okada, R. Okuyama, H. Tagami, S. Aiba, Eosinophilic granulomatous reaction after intradermal injection of hyaluronic acid. Acta Derm-Venereol 88, 69–70 (2008).

59. E. Dayan, C. Chia, A. J. Burns, S. Theodorou, Adjustable Depth Fractional Radiofrequency Combined With Bipolar Radiofrequency: A Minimally Invasive Combination Treatment for Skin Laxity. Aesthet Surg J 39, S112–S119 (2019).

60. B. S. Bloom, J. Emer, D. J. Goldberg, Assessment of safety and efficacy of a bipolar fractionated radiofrequency device in the treatment of photodamaged skin. J Cosmet Laser Ther 14, 208–211 (2012).

61. Z. Chen, J. Y. Seo, Y. K. Kim, S. R. Lee, K. H. Kim, K. H. Cho, H. C. Eun, J. H. Chung, Heat modulation of tropoelastin, fibrillin-1, and matrix metalloproteinase-12 in human skin. J Invest Dermatol 124, 70–78 (2005).

62. J. X. Zhang, Y. He, S. S. Liang, X. Liao, T. Li, Z. Qiao, C. Chang, H. B. Jia, X. W. Chen, Non-invasive, opsin-free mid-infrared modulation activates cortical neurons and accelerates associative learning. Nat Commun 12, (2021).

63. L. Q. Wang, B. Mizaikoff, Application of multivariate data-analysis techniques to biomedical diagnostics based on mid-infrared spectroscopy. Anal Bioanal Chem 391, 1641–1654 (2008).

64. K. J. Wu, C. H. Qi, Z. Zhu, C. L. Wang, B. Song, C. Chang, Terahertz Wave Accelerates DNA Unwinding: A Molecular Dynamics Simulation Study. J Phys Chem Lett 11, 7002–7008 (2020).

65. Y. M. Li, C. Chang, Z. Zhu, L. Sun, C. H. Fan, Terahertz Wave Enhances Permeability of the Voltage-Gated Calcium Channel. J Am Chem Soc 143, 4311–4318 (2021).

66. X. Liu, Z. Qiao, Y. M. Chai, Z. Zhu, K. J. Wu, W. L. Ji, D. G. Li, Y. J. Xiao, L. Q. Mao, C. Chang, Q. Wen, B. Song, Y. S. Shu, Nonthermal and reversible control of neuronal signaling and behavior by midinfrared stimulation. P Natl Acad Sci USA 118, (2021).

67. M. Wang, J. N. Cao, W. K. Amakye, C. C. Gong, Q. Y. Li, J. Y. Ren, Mid infrared light treatment attenuates cognitive decline and alters the gut microbiota community in APP/PS1 mouse model. Biochem Bioph Res Co 523, 60–65 (2020).

